# Overexpression of *Escherichia coli yaiX* Confers Multidrug Resistance and Enhances Virulence in the Silkworm Infection Model

**DOI:** 10.1101/2025.10.16.682852

**Authors:** Kinuka Hongu, Kazuya Ishikawa, Tomoki Kosaki, Shin-Ichi Miyoshi, Kazuyuki Furuta, Chikara Kaito

**Author notes:** Corresponding Author: Chikara Kaito Graduate School of Medicine, Dentistry and Pharmaceutical Sciences Okayama University, Okayama, Japan.

## Abstract

The emergence of bacteria with both antimicrobial resistance and high virulence has become a global health concern, underscoring the urgent need to elucidate the molecular basis underlying these traits. Here, we employed the silkworm (*Bombyx mori*) infection model, which is suitable for high-throughput screening, together with an *Escherichia coli* library containing plasmid clones of all genes from strain W3110, to identify genes whose overexpression enhances virulence. We found that overexpression of the uncharacterized protein YaiX promoted bacterial proliferation in silkworms and increased host lethality. Compared with the empty-vector control, the YaiX-overexpressing strain exhibited resistance to multiple antimicrobial agents with diverse mechanisms of action, including β-lactams, tetracyclines, fluoroquinolones, aminoglycosides, cationic surfactants, and hydrogen peroxide. Sequence analysis revealed that amino acids 18–52 of YaiX contain a transferase hexapeptide domain predicted to form a left-handed parallel β-helix. Overexpression of YaiX mutants lacking regions outside this domain conferred ampicillin resistance, whereas deletion of the hexapeptide domain abolished this phenotype. RNA sequencing and GO enrichment analyses further indicated that YaiX overexpression altered the expression of genes encoding RNA-binding proteins and porins. These findings suggest that YaiX overexpression, through its hexapeptide domain, modulates gene expression and contributes to both multidrug resistance and enhanced virulence in *E. coli*.

## Introduction

The global spread of antimicrobial-resistant bacteria poses a major challenge to the treatment of bacterial infections. *Escherichia coli* is a leading cause of urinary tract infections and a major pathogen responsible for bacteremia and sepsis. Resistant strains have been reported against multiple classes of antibiotics, including β-lactams, aminoglycosides, and fluoroquinolones (1–3). Understanding how the acquisition of antimicrobial resistance affects bacterial pathogenicity is therefore critical for developing strategies to combat resistant infections.

Previous studies have shown that resistance acquisition is often associated with reduced virulence. For example, *E. coli* strains carrying the exogenous *mcr-1* gene acquire colistin resistance through lipid A modification, but this comes at the cost of impaired growth and attenuated virulence in infection models (4). Likewise, in *Staphylococcus aureus*, mutations in the core genome gene *rpoB*, which encodes the RNA polymerase β subunit, confer rifampicin resistance but simultaneously increase susceptibility to oxidative stress and reduce virulence (5–7).

In contrast, recent reports indicate that resistance and virulence can be enhanced simultaneously. In strains such as *Klebsiella pneumoniae* ST23 and *E. coli* ST131, the accumulation of exogenous virulence factors and resistance genes has been shown to promote both traits (8–10). Such bacteria are now recognized in clinical settings as “high-risk pathogens,” characterized by therapeutic intractability and rapid disease progression.

Enhancement of both traits is not limited to exogenous gene acquisition. Alterations in the bacterial core genome can also exert dual effects. For instance, in *Pseudomonas aeruginosa*, mutations in the outer membrane porin OprD confer carbapenem resistance while simultaneously increasing virulence in mice (11–13). In *K. pneumoniae*, deletion of the MarR-family repressor RamR not only confers resistance to colistin and the antimicrobial peptide LL-37 but also enhances evasion of macrophage phagocytosis and virulence in murine models (14).

We have previously reported, using the silkworm infection model—which is advantageous for exploratory studies due to its low cost and minimal ethical concerns (15, 16)—that mutations or deletions in *E. coli* core-genome genes can simultaneously increase both pathogenicity and resistance. Specifically, amino acid substitutions in the LPS transporters LptD and LptE, deletion of the periplasmic polysaccharide-synthesizing enzymes OpgG and OpgH, and loss of the outer membrane lipoprotein MlaA, a component of the phospholipid transport system, all enhanced virulence toward silkworms and conferred resistance to multiple antibiotics (17–19). In *Bacillus subtilis*, disruption of the putative glycosyltransferase YkcB was also shown to increase both pathogenicity and vancomycin resistance (20). These findings suggest the presence of intrinsic systems by which core-genome alterations can enhance both virulence and antimicrobial resistance, independent of exogenous gene acquisition. However, the mechanisms underlying such intrinsic dual enhancement remain poorly understood.

In this study, we performed a genome-wide analysis of *E. coli* using the silkworm infection model to identify genes whose overexpression enhances pathogenicity. Our results demonstrate that overexpression of the previously uncharacterized gene *yaiX* increases virulence in silkworms while conferring resistance to multiple antibiotics.

## Results

### Overexpression of *yaiX* confers a survival advantage and enhances lethality in silkworms

We screened for genes whose overexpression conferred a survival advantage in silkworms by passaging the ASKA library—*E. coli* strains individually overexpressing each gene—in the silkworm infection model (**Fig. 1A; Table S1**). Colonies recovered from dead silkworms were analyzed by colony PCR, and the occupancy ratio of each identified clone within an ASKA library plate was calculated. Strains with an occupancy ratio exceeding 25% were subjected to 1:1 competition assays against the empty-vector control strain (EV) in silkworms. Among these, 26 strains exhibited occupancy ratios greater than 50%, indicating a survival advantage over the EV strain (**Table S1**). Notably, four strains reached 100% occupancy, demonstrating a strong survival advantage during silkworm infection. Of these, the *yaiX*-overexpressing strain, encoding a previously uncharacterized protein, was selected for further characterization.

**Figure 1.**
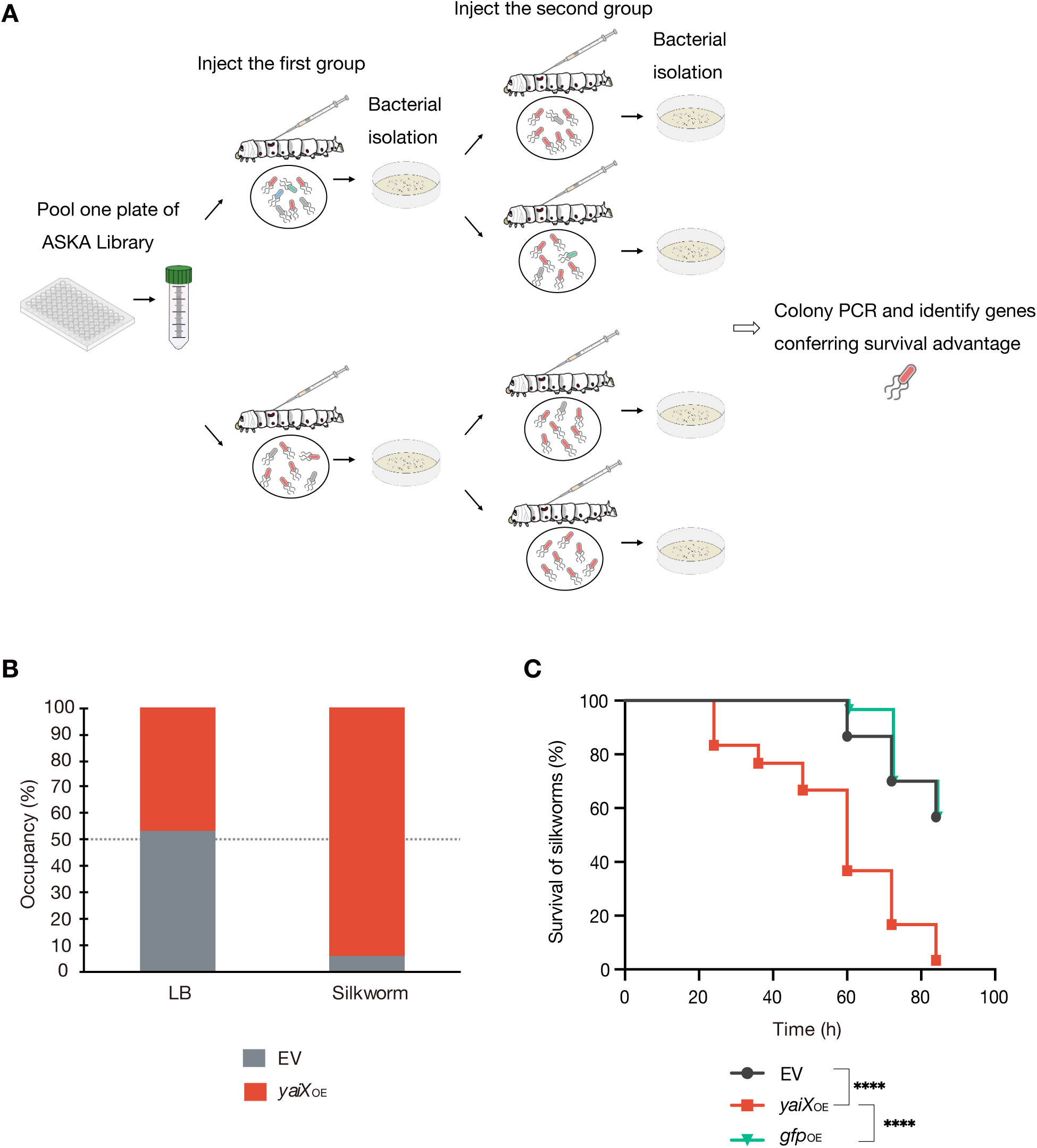
Overexpression of *yaiX* confers a survival advantage and enhances killing activity in silkworms. A. Schematic of the silkworm survival screening. Each 96-well plate of the ASKA library was cultured overnight in LB medium containing 0.1 mM IPTG, and pooled cultures were injected into two silkworms. Immediately after host death, hemolymph was collected and bacteria were recovered. The recovered bacteria were re-infected into silkworms, and bacteria were again isolated from hemolymph after host death and plated on LB agar containing chloramphenicol. Five colonies per silkworm (20 colonies per ASKA plate) were randomly selected, and the overexpressed genes carried on the plasmids were identified by PCR and Sanger sequencing to calculate the occupancy rate. In total, 56 ASKA plates were analyzed; strains with an occupancy rate >25% were designated as having a survival advantage in silkworms. B. 1:1 competition assays of EV and *yaiX*-overexpressing strains in LB broth and in silkworms. For the LB broth assay, a 1:1 mixture (100 µL, 5.43 × 10⁴ CFU) of overnight cultures of EV and *yaiX*-overexpressing strains was inoculated into 5 mL fresh LB and cultured overnight at 37 °C. Bacteria were plated, and occupancy was determined by colony PCR. For the silkworm assay, a 1:1 mixture (50 µL, 4.83 × 10³ CFU) was injected into silkworms; after host death, bacteria were recovered from hemolymph and occupancy was determined by colony PCR. Data represent the mean of three independent experiments. C. Survival of silkworms injected with EV, *yaiX*-overexpressing, or *gfp*-overexpressing strains. Suspensions were adjusted to OD_600_ = 7.0 (3.80 × 10⁶ CFU/mL), and 50 µL was injected (n = 10 per group). Data were pooled from three independent experiments (n = 30). Statistical analysis used the log-rank test. **** P < 0.0001.

To determine whether the survival advantage of the *yaiX*-overexpressing strain was specific to infection, we performed competition assays both in LB broth and during silkworm infection. In LB broth, the occupancy rates of the EV and *yaiX*-overexpressing strains were 53% and 47%, respectively. In contrast, during silkworm infection, occupancy was 6% for the EV strain and 94% for the *yaiX*-overexpressing strain (**Fig. 1B**). These results indicate that the advantage conferred by *yaiX* overexpression is specific to the infection context.

We next tested whether this survival advantage also increased killing activity in silkworms. EV, GFP-overexpressing, and *yaiX*-overexpressing strains were injected into silkworm hemolymph, and survival was monitored every 12 h for up to 84 h. Silkworms injected with EV or GFP-overexpressing strains maintained >50% survival at 84 h. In contrast, >50% of silkworms injected with the *yaiX*-overexpressing strain were dead by 60 h, and nearly all were dead by 84 h (**Fig. 1C**). These data demonstrate that *yaiX* overexpression enhances the killing activity of *E. coli* in the silkworm model.

### Overexpression of *yaiX* confers resistance to diverse antimicrobial agents, including hydrogen peroxide

Given that *yaiX* overexpression enhanced virulence in silkworms, we hypothesized that the strain might acquire resistance to oxidative stress, a key innate immune mechanism. EV, GFP-overexpressing, and *yaiX*-overexpressing strains were inoculated into medium containing 18 mM H₂O₂ at equal optical densities, and growth was monitored for 12 h. In the presence of 18 mM H₂O₂, neither the EV nor GFP-overexpressing strains grew, whereas the *yaiX*-overexpressing strain exhibited robust growth (**Fig. 2A**).

**Figure 2.**
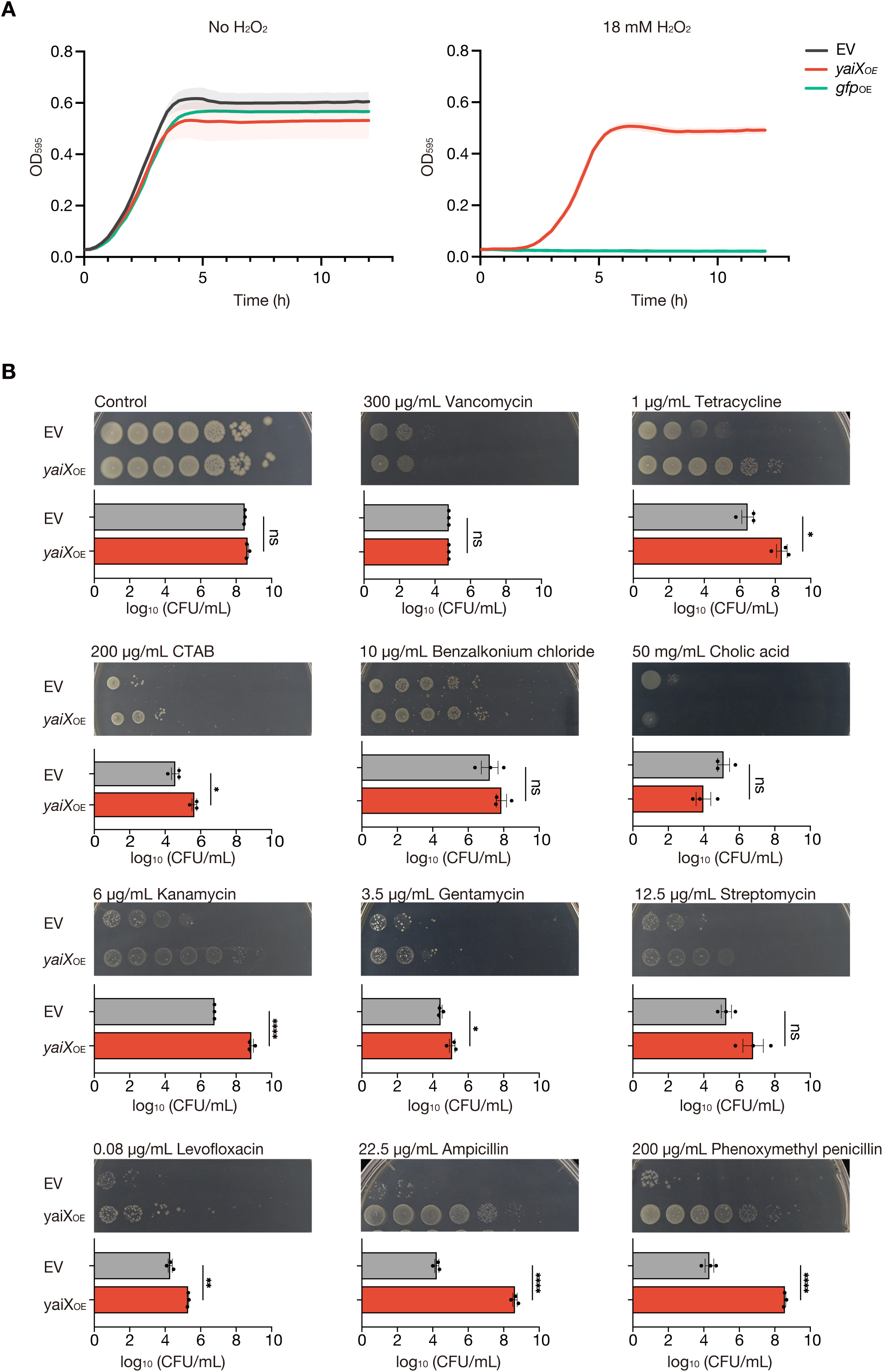
Overexpression of *yaiX* confers multidrug resistance and hydrogen peroxide resistance. A. Hydrogen peroxide resistance of the *yaiX*-overexpressing strain. Overnight cultures of EV, *yaiX*-overexpressing, or *gfp*-overexpressing strains were inoculated 1:100 into LB broth containing 0.1 mM IPTG with or without 18 mM H₂O₂ and incubated at 37 °C for 12 h. The y-axis shows OD_595_; the x-axis shows time. Data are mean ± SEM from three biological replicates. B. Multidrug resistance of the *yaiX*-overexpressing strain. Overnight cultures of EV and *yaiX*-overexpressing strains were adjusted to OD_600_ = 1.0, serially diluted 10-fold, and spotted onto LB agar supplemented with the indicated antimicrobial agents and 0.1 mM IPTG. Plates were incubated at 37 °C. Representative images from three independent experiments are shown; graphs display mean ± SEM from ≥3 independent experiments. Statistical analysis used an unpaired t-test. ns, P > 0.5; P < 0.01; *P < 0.001; **P < 0.001; **** P < 0.0001.

We next examined whether *yaiX* overexpression also conferred resistance to other antimicrobial agents. Spot assays revealed that, compared with the EV strain, the *yaiX*-overexpressing strain maintained higher CFU counts in the presence of tetracycline, cetyltrimethylammonium bromide (CTAB), kanamycin, gentamicin, levofloxacin, ampicillin, and phenoxymethylpenicillin (**Fig. 2B**). In contrast, there were no significant differences between the EV strain and the *yaiX*-overexpressing strain in the presence of vancomycin, benzalkonium chloride, cholic acid, or streptomycin (**Fig. 2B**). These findings indicate that *yaiX* overexpression confers broad resistance to antimicrobial agents with diverse mechanisms of action, including β-lactams and aminoglycosides.

### RNA-seq analysis of the *yaiX*-overexpressing strain

To further investigate *yaiX* function, we performed RNA-seq analysis. Compared with the EV strain, 17 genes were significantly upregulated and 7 genes were significantly downregulated in the *yaiX*-overexpressing strain (**Fig. 3A**). GO enrichment analysis of 39 genes with altered expression (p < 0.1) revealed enrichment in categories related to RNA-binding functions and porin activity (**Fig. 3B**).

**Figure 3.**
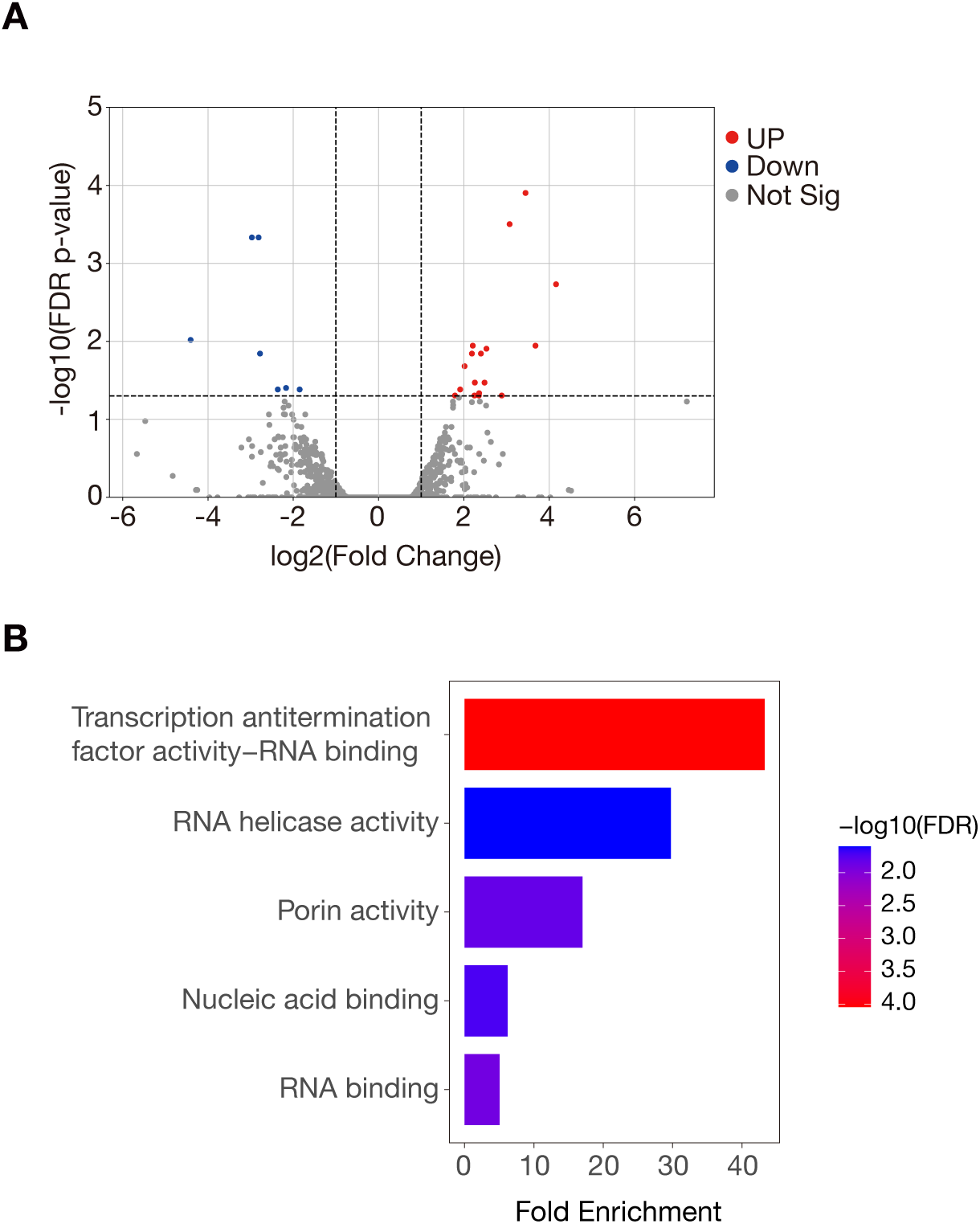
RNA-seq analysis of the *yaiX*-overexpressing strain. A. Volcano plot showing differences in gene expression between EV and *yaiX*-overexpressing strains. The x-axis shows log₂ fold change; the y-axis shows −log₁₀ p-value. Significantly altered genes (|log₂ fold change| > 1 and p < 0.05) are shown in red (upregulated) or blue (downregulated); others are in gray. B. GO enrichment analysis of genes differentially expressed between EV and *yaiX*-overexpressing strains with p < 0.1. Analysis was performed using ShinyGO v0.80.

### Contribution of differentially expressed genes to antimicrobial resistance and virulence

To determine whether the differentially expressed genes contribute to *yaiX* function, we examined their roles in ampicillin resistance. Overexpression of the 17 upregulated genes did not confer resistance (**Fig. 4A**). In contrast, among the 7 downregulated genes, the *ompF* deletion mutant exhibited strong ampicillin resistance (**Fig. 4B**), consistent with previous reports (21, 22).

**Figure 4.**
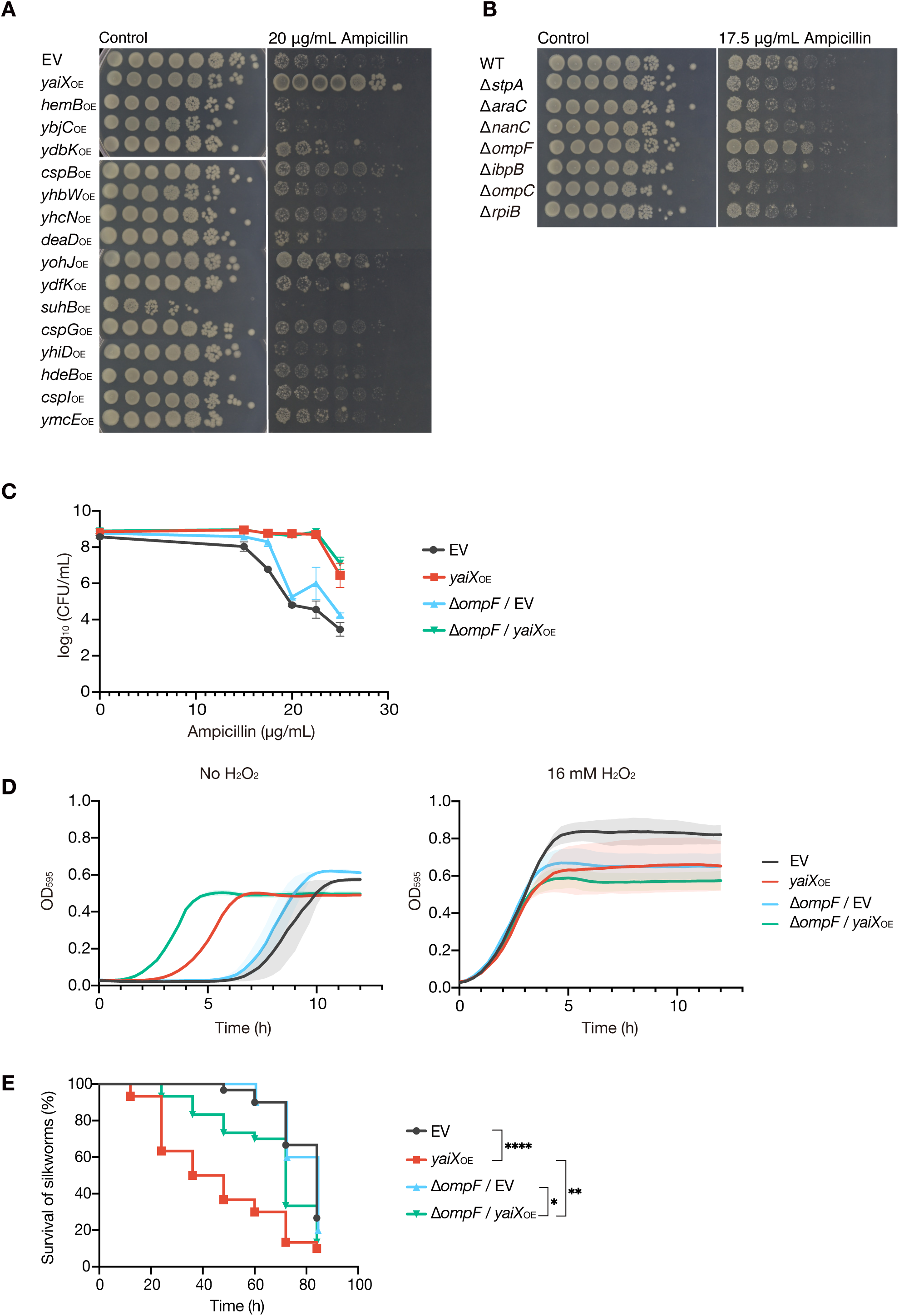
Effects of differentially expressed genes in the *yaiX*-overexpressing strain on antibiotic resistance and virulence. A. Effect of genes upregulated (p < 0.05) in the *yaiX*-overexpressing strain on ampicillin resistance. Overexpression strains were cultured overnight in medium containing 0.1 mM IPTG, adjusted to OD_600_ = 1.0, and spotted onto LB agar containing 0.1 mM IPTG with or without ampicillin (20 μg/mL); plates were incubated at 37 °C. B. Effect of genes downregulated (p < 0.05) in the *yaiX*-overexpressing strain on ampicillin resistance. Deletion mutants were cultured overnight, adjusted to OD_600_ = 1.0, and spotted onto LB agar containing 0.1 mM IPTG with or without ampicillin (17.5 μg/mL); plates were incubated at 37 °C. C. Ampicillin resistance of the *E. coli* parent strain carrying either the EV or *yaiX*-overexpression plasmid, and the Δ*ompF* strain carrying either the EV or *yaiX*-overexpression plasmid. Overnight cultures were adjusted to OD_600_ = 1.0 and spotted onto LB agar containing 0.1 mM IPTG, with or without ampicillin (15–25 μg/mL). Data are mean ± SEM of three independent experiments. D. Hydrogen peroxide resistance of the *E. coli* parent strain carrying either the EV or *yaiX*-overexpression plasmid, and the Δ*ompF* strain carrying either the EV or *yaiX*-overexpression plasmid. Strains were cultured in LB broth containing 0.1 mM IPTG with or without 16 mM H₂O₂. The y-axis shows OD_595_; the x-axis shows time. Data are mean ± SEM of three biological replicates. E. Virulence of the *E. coli* parent strain carrying either the EV or *yaiX*-overexpression plasmid, and the Δ*ompF* strain carrying either the EV or *yaiX*-overexpression plasmid in silkworms. Overnight cultures were adjusted to OD_600_ = 7.0, and 10 silkworms were injected per strain. Data were pooled from three independent experiments (n = 30). Statistical analysis used the log-rank test.

We therefore tested *yaiX* overexpression in a Δ*ompF* background. The *yaiX*-overexpressing Δ*ompF* strain showed greater ampicillin resistance than the Δ*ompF* strain carrying EV (**Fig. 4C**). Similarly, the *yaiX*-overexpressing Δ*ompF* strain displayed increased resistance to H₂O₂ compared with the Δ*ompF* strain carrying EV (**Fig. 4D**). These results indicate that *yaiX* overexpression confers resistance to both ampicillin and hydrogen peroxide independently of *ompF*.

Finally, we assessed virulence in silkworms. The *yaiX*-overexpressing Δ*ompF* strain killed silkworms more rapidly than the Δ*ompF* strain carrying EV (**Fig. 4E**). However, the degree of enhancement was smaller than that observed in the wild-type background (**Fig. 4E**). Thus, *yaiX* overexpression enhances virulence in a manner partially dependent on *ompF*.

### The hexapeptide domain is essential for YaiX activity

YaiX is a 71-amino acid protein predicted to function as an acyltransferase and annotated as a pseudogene disrupted by an insertion sequence (23, 24). Structural prediction using AlphaFold2 revealed that its central region contains a hexapeptide domain forming a β-helix structure (**Fig. 5A**).

**Figure 5.**
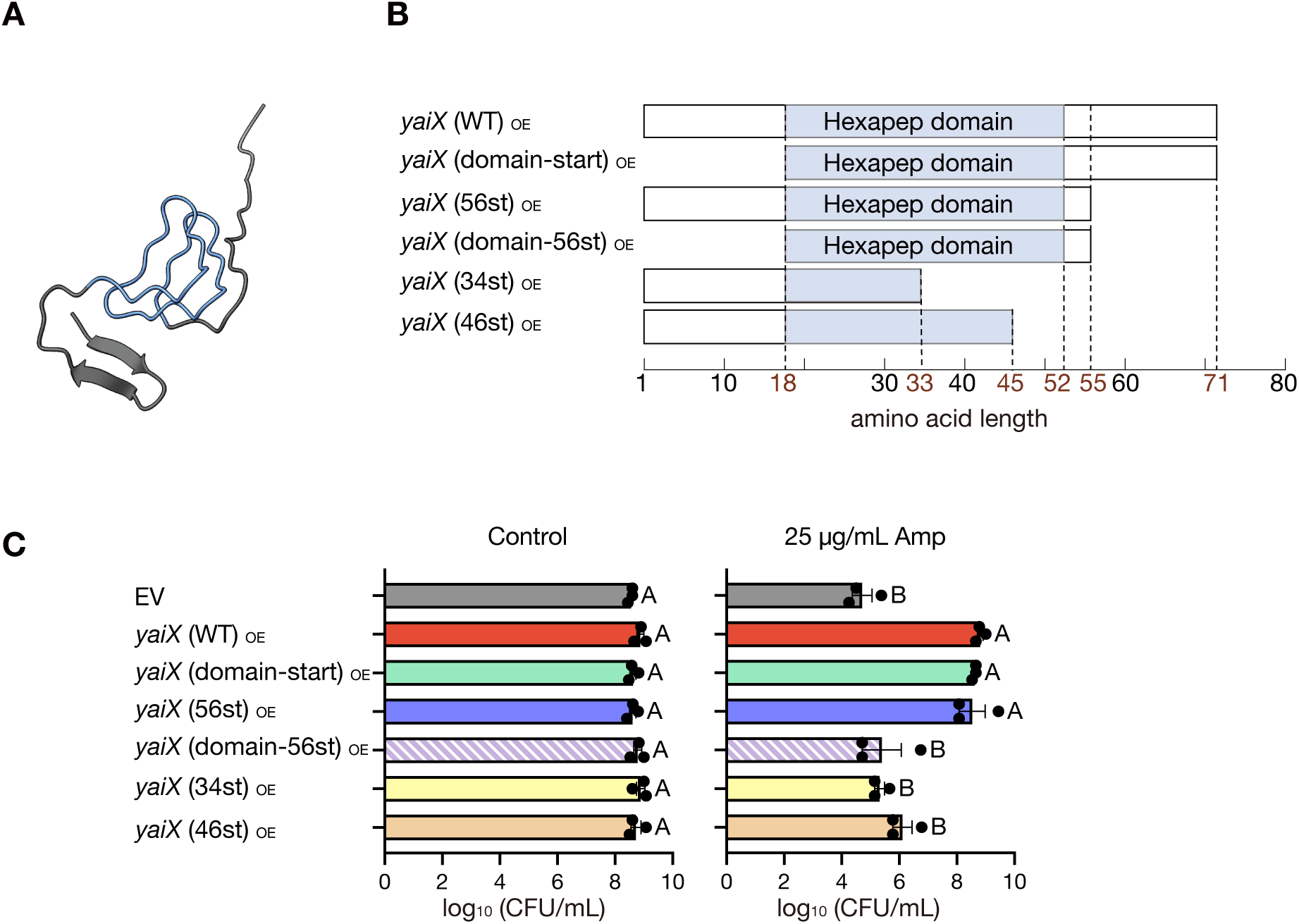
The hexapeptide domain is essential for YaiX activity. A. Predicted 3D structure of YaiX generated from its amino acid sequence using AlphaFold2. B. Schematic of YaiX deletion mutants used in this study. The blue region indicates the hexapeptide domain; numbers denote amino-acid residue positions. C. Ampicillin resistance of overexpression strains carrying the YaiX deletion mutants shown in panel D. Overnight cultures of the *E. coli* parent strain carrying either the EV, the *yaiX*-overexpression plasmid, or YaiX deletion mutant-overexpression plasmids were adjusted to OD_600_ = 1.0 and spotted onto LB agar containing 0.1 mM IPTG, with or without ampicillin (25 μg/mL). Plates were incubated at 37 °C. Data are mean ± SEM of three independent experiments. Statistical analysis used one-way ANOVA followed by Tukey’s multiple-comparison test.

To test the role of this domain, we constructed plasmids expressing YaiX deletion variants (**Fig. 5B**). Strains expressing YaiX(domain-start), which lacked the N-terminal flanking region of the domain, or YaiX(56st), which lacked the C-terminal flanking region of the domain, conferred strong ampicillin resistance comparable to that of wild-type YaiX (**Fig. 5C**). In contrast, YaiX(domain-56st), which lacked both flanking regions of the domain, failed to confer resistance. Likewise, strains expressing YaiX variants with partial deletions within the domain combined with deletion of the C-terminal flanking region (YaiX-34st or YaiX-46st) also lost resistance (**Fig. 5C**). These findings demonstrate that the hexapeptide domain is essential for YaiX-mediated ampicillin resistance.

### The *yaiX* sequence is conserved in more than half of *E. coli* strains

We analyzed the presence and distribution of *yaiX* across multiple *E. coli* strains. Phylogenetic analysis of 58 genomes from KEGG revealed three groups: strains with the 71-aa variant, strains with the 236-aa variant, and strains lacking *yaiX* (**Fig. 6A, 6B**).

**Figure 6.**
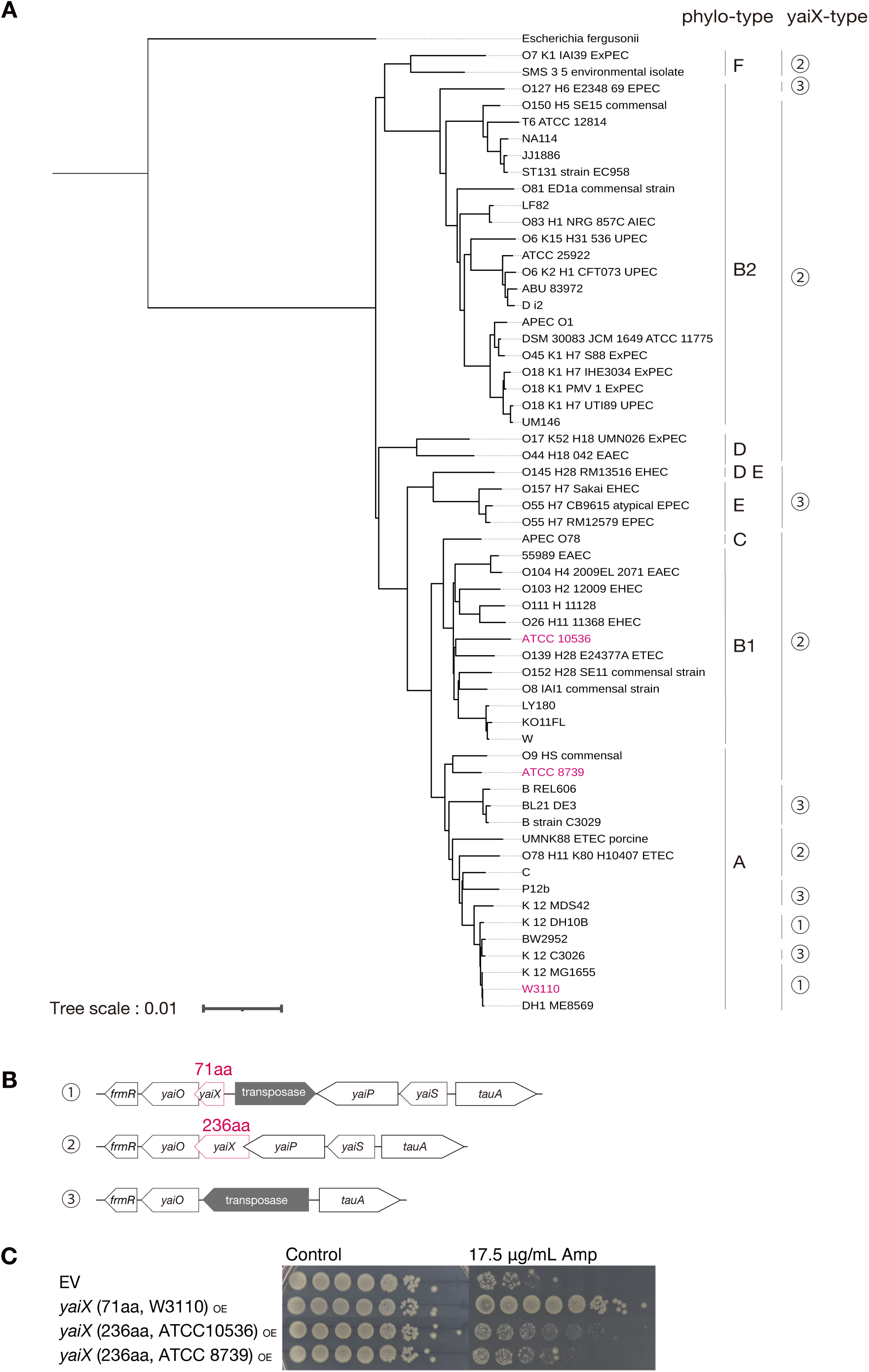
The *yaiX* sequence is conserved in more than half of *E. coli* strains. A. Phylogenetic tree of 58 *E. coli* strains. Whole-genome sequences were downloaded from KEGG, pairwise genomic distances were calculated using Mash, and the tree was inferred using FastME. The Newick-format tree was visualized with iTOL. *Escherichia fergusonii* was used as the outgroup. B. Gene organization surrounding *yaiX* in *E. coli*. Numbers 1–3 correspond to the strains shown in panel A. The *yaiX* locus and its amino-acid length are indicated in red. C. Ampicillin resistance of EV, 71-aa *yaiX*-overexpressing, and 236-aa *yaiX*-overexpressing strains. Overnight cultures were adjusted to OD_600_ = 1.0 and spotted onto LB agar containing 0.1 mM IPTG with or without ampicillin (17.5 μg/mL). Plates were incubated overnight at 37 °C.

Functional assays showed that the 236-aa variant conferred ampicillin resistance compared with EV, but the level of resistance was weaker than that conferred by the 71-aa variant (**Fig. 6C**). These data indicate that *yaiX* is conserved in many *E. coli* strains and that the shorter 71-aa form provides stronger resistance.

## Discussion

In this study, we demonstrated that overexpression of *yaiX* simultaneously enhanced virulence and multidrug resistance in *E. coli*. To our knowledge, this is the first report showing that overexpression of a single intrinsic gene can confer both traits.

Overexpression of *yaiX* conferred resistance to multiple antimicrobial agents, including β-lactams, aminoglycosides, levofloxacin, tetracycline, CTAB, and H₂O₂. As reactive oxygen species, including H₂O₂, are key antimicrobial effectors of the innate immune system in silkworms (5, 25, 26), resistance to H₂O₂ likely contributes to the increased bacterial burden and lethality observed in infected silkworms. Since these antimicrobial agents act on distinct bacterial targets, *yaiX* likely does not interfere directly with each mechanism but may instead affect a common trait such as membrane permeability.

RNA-seq analysis showed altered expression of genes encoding RNA-binding proteins and porins. Downregulation of *ompF*, a major channel for β-lactams (21, 22), was observed, and Δ*ompF* strains confirmed its role in resistance. However, *yaiX* overexpression still conferred ampicillin resistance and virulence in the Δ*ompF* background, indicating both *ompF*-dependent and -independent pathways. None of the other differentially expressed genes contributed to resistance, highlighting the need to elucidate *ompF*-independent mechanisms.

YaiX is a 71-amino-acid protein containing a hexapeptide domain (23), whose function has not yet been characterized. We found that deletion of the hexapeptide domain abolished *yaiX*-mediated resistance, suggesting that the domain is critical for its activity. Although this domain is conserved among various transferases, the substrate of YaiX remains unknown. Future studies should test whether purified YaiX has acyltransferase activity.

Comparative genomics revealed that *yaiX* is conserved in both 71-aa and 236-aa forms, with the shorter variant conferring stronger resistance. This suggests that pseudogenization-driven truncation may provide a selective advantage by enhancing resistance and virulence. Notably, no Shine– Dalgarno sequence was identified upstream of *yaiX* in either variant, suggesting minimal expression under normal conditions. Genomic alterations, such as insertion sequences, may activate expression under certain conditions.

Finally, our findings highlight both commonalities and differences between *yaiX*-overexpressing strain and previously described mutants (LptD, LptE, OpgG, OpgH, and MlaA) (17–19). Whereas those mutants conferred vancomycin resistance, *yaiX* overexpression did not. However, resistance to levofloxacin was shared by *yaiX*-overexpressing strain and several of these mutants. These distinct susceptibility profiles indicate that multiple, mechanistically diverse pathways can simultaneously enhance pathogenicity and antimicrobial resistance.

Future studies aimed at clarifying the molecular mechanisms underlying YaiX activity and comparing it with other intrinsic resistance–virulence determinants will provide important insights into the evolution of high-risk *E. coli* strains.

## Materials and Methods

### Bacterial strains and culture conditions

*Escherichia coli* BW25113, AG1, ATCC10536, and ATCC8739 were cultured aerobically at 37 °C in LB medium. Mutant strains carrying a kanamycin resistance gene were grown in LB medium supplemented with kanamycin (100 μg/mL). Strains harboring the pCA24N vector or gene-overexpression plasmids were maintained in LB medium supplemented with chloramphenicol (30 μg/mL). For gene overexpression, strains were cultured in the presence of 0.1 mM IPTG. Details of the strains and plasmids used in this study are provided in Tables 1 and 2.

**Table 1.**
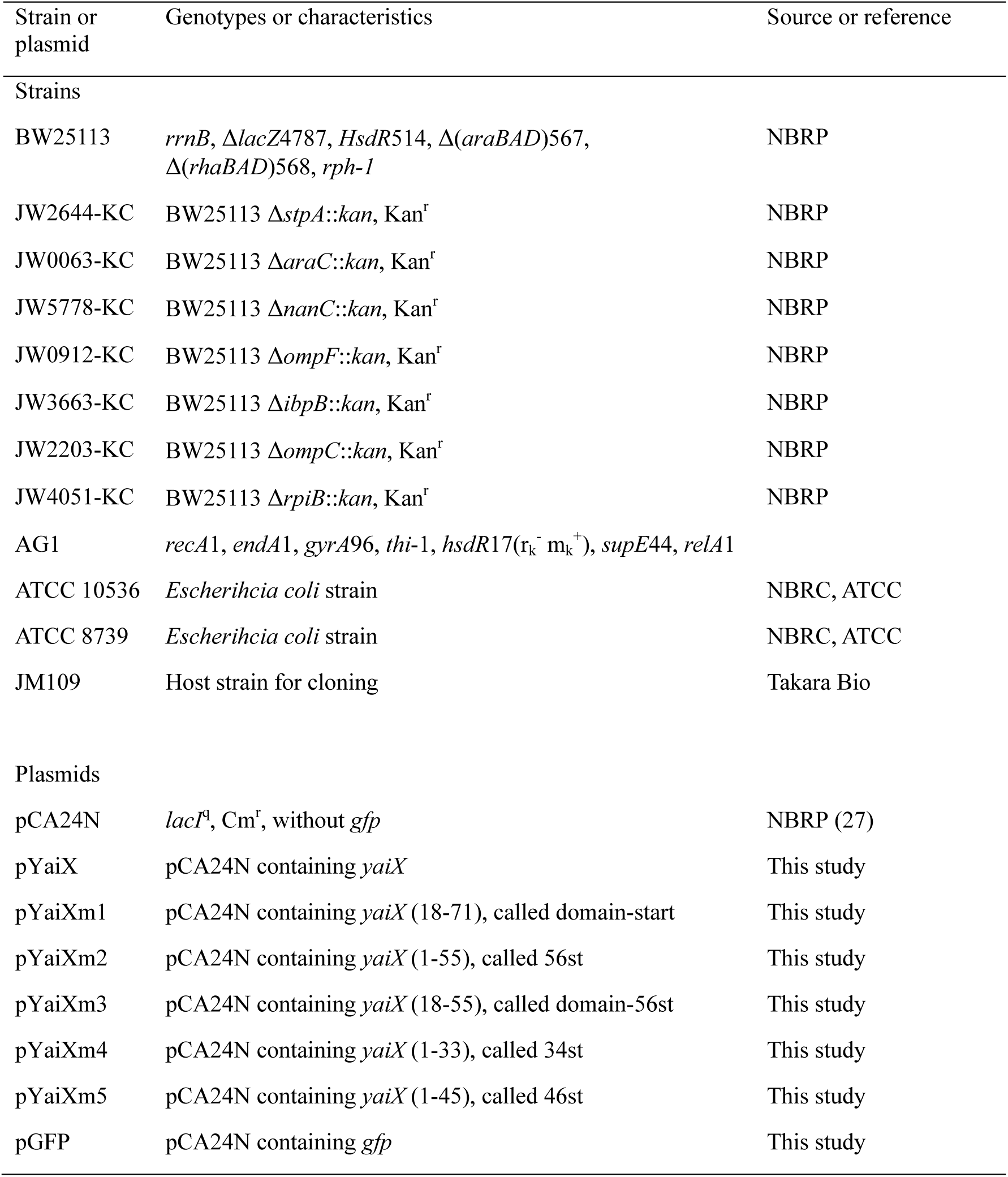
List of bacterial strains and plasmids used.

| Strain or plasmid | Genotypes or characteristics | Source or reference |
| --- | --- | --- |
| Strains |  |  |
| BW25113 | *rrnB*, Δ*lacZ*4787, *HsdR*514, $\Delta$(*araBAD*)567, $\Delta$(*rhaBAD*)568, *rph-1* | NBRP |
| JW2644-KC | BW25113 Δ*stpA*::*kan*, Kan<sup>r</sup> | NBRP |
| JW0063-KC | BW25113 Δ*araC*::*kan*, Kan<sup>r</sup> | NBRP |
| JW5778-KC | BW25113 Δ*nanC*::*kan*, Kan<sup>r</sup> | NBRP |
| JW0912-KC | BW25113 Δ*ompF*::*kan*, Kan<sup>r</sup> | NBRP |
| JW3663-KC | BW25113 Δ*ibpB*::*kan*, Kan<sup>r</sup> | NBRP |
| JW2203-KC | BW25113 Δ*ompC*::*kan*, Kan<sup>r</sup> | NBRP |
| JW4051-KC | BW25113 Δ*rpiB*::*kan*, Kan<sup>r</sup> | NBRP |
| AG1 | *recA1*, *endA1*, *gyrA96*, *thi-1*, *hsdR17*(r<sub>k</sub><sup>-</sup> m<sub>k</sub><sup>+</sup>), *supE44*, *relA1* |  |
| ATCC 10536 | *Escherihcia coli* strain | NBRC, ATCC |
| ATCC 8739 | *Escherihcia coli* strain | NBRC, ATCC |
| JM109 | Host strain for cloning | Takara Bio |
| Plasmids |  |  |
| pCA24N | *lacI*<sup>q</sup>, Cm<sup>r</sup>, without *gfp* | NBRP (27) |
| pYaiX | pCA24N containing *yaiX* | This study |
| pYaiXm1 | pCA24N containing *yaiX* (18-71), called domain-start | This study |
| pYaiXm2 | pCA24N containing *yaiX* (1-55), called 56st | This study |
| pYaiXm3 | pCA24N containing *yaiX* (18-55), called domain-56st | This study |
| pYaiXm4 | pCA24N containing *yaiX* (1-33), called 34st | This study |
| pYaiXm5 | pCA24N containing *yaiX* (1-45), called 46st | This study |
| pGFP | pCA24N containing *gfp* | This study |

**Table 2.**
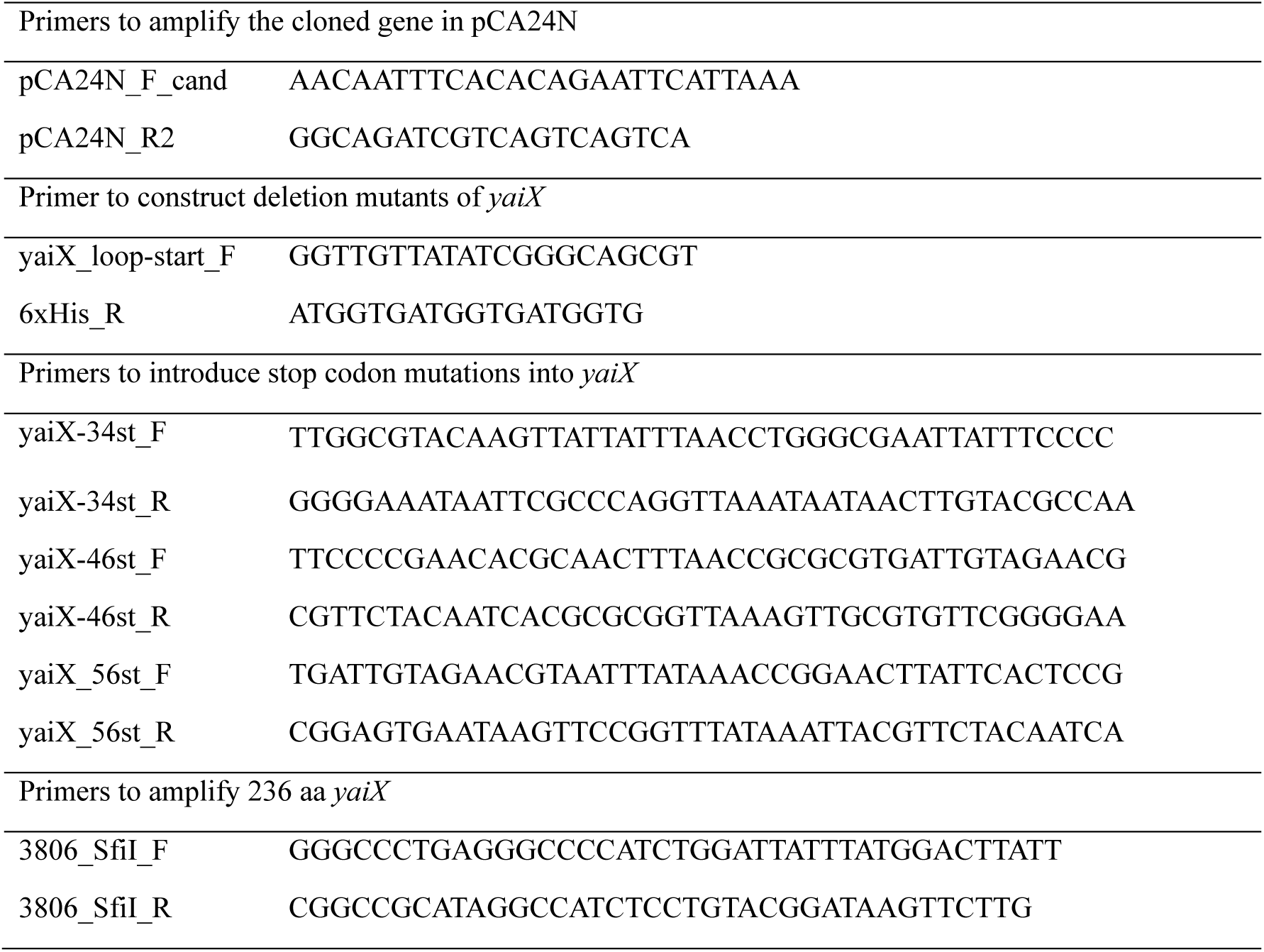
Primers used in this study.

### Construction of gene-overexpression strains

Gene-overexpression strains were generated by isolating the corresponding pCA24N plasmids from the ASKA library (27) and introducing them into *E. coli* BW25113. For the 236-amino-acid form of *yaiX* derived from strains ATCC10536 and ATCC8739, the genes were PCR-amplified using primers listed in Table 2 and cloned into the SfiI restriction site of pCA24N. The resulting plasmids were transformed into *E. coli* BW25113.

To construct pYaiXm1 carrying a deletion mutant of *yaiX*, DNA fragments were amplified by inverse PCR using the primers listed in Table 2 with pYaiX as a template, and the products were self-ligated. To construct pYaiXm2, pYaiXm3, pYaiXm4, and pYaiXm5 carrying *yaiX* genes with introduced stop codons, thermal cycling was performed as previously described (28). Briefly, thermal cycling was carried out using the primers listed in Table 2 with pYaiX as a template. The mutations were confirmed by Sanger sequencing, and the resulting plasmids were introduced into *E. coli* BW25113 by electroporation.

### Gene deletion strains

Gene deletion strains were obtained from the Keio collection (29). For *ompF* deletion, the mutation was transferred from the original JW0912-KC strain into *E. coli* BW25113 by phage P1 transduction, and the mutant was reconstructed. This newly generated Δ*ompF* strain was then used to evaluate the effects of *yaiX* overexpression.

### In vivo survival screening in silkworms

Each 96-well plate of the ASKA library was first cultured overnight in LB medium containing 30 μg/mL chloramphenicol. Cultures were transferred into fresh LB medium (100 µL/well) supplemented with 0.1 mM IPTG and 30 μg/mL chloramphenicol in a 96-well plate and incubated overnight. Bacterial suspensions from all wells were pooled, resuspended in NaCl buffer (0.9% NaCl, 0.1 mM IPTG, 30 μg/mL chloramphenicol), and adjusted to an OD_600_ of 0.18. This suspension was injected into two silkworms. After host death, hemolymph was collected and plated on LB agar containing 30 μg/mL chloramphenicol. Colonies were resuspended in NaCl buffer, adjusted to the same OD_600_, and injected into two additional silkworms. Bacteria were again recovered after host death. Five colonies were randomly selected from each silkworm (20 colonies per plate) and analyzed by PCR and Sanger sequencing to identify the overexpressed genes. Occupancy rates were calculated, and strains with an occupancy rate >25% were considered to have a survival advantage. A total of 56 plates were screened.

For candidate strains, the corresponding pCA24N plasmids were isolated from the ASKA library and introduced into *E. coli* BW25113. In parallel, the empty pCA24N vector (without GFP) was transformed into BW25113 as a control. Each overexpression strain and the EV strain were adjusted to the same OD_600_, mixed at a 1:1 ratio, and 50 μL of the mixture was injected into three silkworms. Silkworms were incubated at 37 °C until death. Hemolymph was collected immediately after death and plated on LB agar containing 30 μg/mL chloramphenicol. Four colonies from each silkworm (12 colonies total) were analyzed by PCR, and occupancy rates were determined.

### Silkworm killing assay

*E. coli* BW25113 strains carrying either the EV or gene-overexpression plasmids were cultured overnight at 37 °C in LB medium containing 0.1 mM IPTG and 30 μg/mL chloramphenicol. Cultures were centrifuged at 4,050 × g for 10 min, and pellets were resuspended in NaCl buffer and adjusted to an OD_600_ of 7.0. Aliquots of 50 μL were injected into 10 silkworms per strain. Silkworms were incubated at 37 °C, and survival was monitored every 12 h. Viability was assessed by the presence or absence of a response to mechanical stimulation.

### Assessment of resistance to antimicrobial agents

LB agar was autoclaved and supplemented with 0.1 mM IPTG, 30 μg/mL chloramphenicol, and the indicated antimicrobial agents, then poured into Petri dishes. Overnight cultures of *E. coli* were serially diluted 10-fold in 96-well microplates, and 5 µL of each dilution was spotted onto the agar plates. Plates were incubated overnight at 37 °C, and colony formation was recorded using a digital camera. Colony counts were performed as described previously (30).

### Growth in the presence of hydrogen peroxide

*E. coli* BW25113 strains were cultured overnight at 37 °C in LB medium containing 0.1 mM IPTG and 30 μg/mL chloramphenicol. Cultures were adjusted to the same OD_600_ and exposed to hydrogen peroxide in LB broth supplemented with 0.1 mM IPTG and 30 μg/mL chloramphenicol. Growth was monitored at 37 °C for 12 h by measuring OD_595_ using a microplate reader.

### RNA sequencing

Total RNA was extracted with minor modifications to a published method (31). Overnight cultures of the EV and *yaiX*-overexpressing (*yaiX*_OE_) strains (50 µL each) were inoculated into 5 mL LB medium and grown aerobically at 37 °C. At OD_600_ = 0.7, 1.8 mL of culture was mixed with 200 µL of 5% ethanol-saturated phenol, cooled on ice for 5 min, and centrifuged at 21,500 × g for 2 min. Pellets were frozen in liquid nitrogen and stored at −80 °C for 2 h, then resuspended in 200 µL lysis buffer (TE buffer containing 1% lysozyme and 1% SDS) and incubated at 65 °C for 2 min. RNA was purified using the RNeasy Mini Kit (Qiagen). Ribosomal RNA was removed with the NEBNext rRNA Depletion Kit (NEB), and libraries were prepared using the TruSeq Stranded Total RNA Kit (Illumina). Sequencing was performed on a NovaSeq 6000 system (Illumina), generating ≥4 Gbp of 100-bp paired-end reads per sample. Data were analyzed using CLC Genomics Workbench v11.0. Reads were mapped to the *E. coli* W3110 reference genome (NCBI RefSeq NC_007779.1), and RPKM values were compared between the EV and *yaiX*_OE_ strains. Experiments were performed independently in duplicate. GO analysis was conducted with ShinyGO 0.80 (https://bioinformatics.sdstate.edu/go80/) (32).

### Phylogenetic analysis based on whole-genome distances

Whole-genome sequences (FASTA format) of *E. coli* strains were obtained from KEGG. Pairwise genomic distances were calculated with Mash (v2.2.2) (33). Outputs were converted to a PHYLIP-format distance matrix using an in-house script, and distance-based phylogenetic trees were constructed with FastME (v2.1.6.3) using default parameters (34). Trees were visualized with iTOL (Interactive Tree of Life; v7) (35), with *Escherichia fergusonii* FDAARGOS_1499 designated as the outgroup.

### Statistical analysis

Data were analyzed using GraphPad Prism v10. Statistical tests are specified in the corresponding figure legends.

## Acknowledgement

This study was supported by Japan Society for the Promotion of Science (JSPS) Grants-in-Aid for Scientific Research (Grants 23K24131, 23K06130, 24K01760) and the Program of the Japan Initiative for Global Research Network&Link on Infectious Diseases (J-GRID+), JP24wm0125004, from Ministry of Education, Culture, Sports, Science and Technology in Japan (MEXT), and Japan Agency for Medical Research and Development (AMED). We thank the National BioResource Project-E. coli (National Institute of Genetics, Japan) for providing the E. coli Keio collection.

